# Shift in bacterial taxa precedes morphological plasticity in a larval echinoid

**DOI:** 10.1101/640953

**Authors:** Tyler J. Carrier, Adam M. Reitzel

## Abstract

Morphological plasticity is an adaptive response to heterogenous environments when a fitness advantage is conferred. Larval sea urchins, for example, increase individual fitness in dilute feeding environments by elongating their feeding structure. Morphological plasticity for larval sea urchins is also coupled with significant shifts in the associated bacterial community, but whether this response occurs before, during, or following the expression of plasticity is unclear. Using the sea urchin *Lytechinus variegatus*, we define the temporal pattern of the associated bacterial community throughout the expression of morphological plasticity. From prefeeding through plasticity, we observed that *L. variegatus* larvae exhibit a four-stage successional pattern and the relatedness of the larval-associated bacterial community directly reflects morphological plasticity and does so prior to expression of the environmental-specific morphology. Based on the structure of the larval-associated bacterial communities, the expression of morphological plasticity correlates short-arm larvae deviating from the microbial trajectory of pre-plastic siblings. Taken together, these data suggest that a holobiont may exhibit shifts in the associated bacterial community corresponding with the environmental variation in absence or anticipation of morphological plasticity.

## 1. Introduction

As the environment ebbs and flows, animals from diverse phylogenetic lineages may modify phenotypic traits to better match environmental conditions [1–3]. When phenotypic plasticity—the potential to produce a range of phenotypes from a single genotype—is induced and confers a fitness advantage, the response is considered adaptive and an evolvable trait influenced by natural selection [1–3]. For morphological characters to exhibit plasticity favored by selection, abiotic or biotic stimuli must be reliable and detected at an appropriate time scale to elicit a developmental change in a structure that results in a phenotype-environment match [1, 4, 5]. Plasticity in morphology and other life history characters are, thus, viewed as a product of host genotype and the environment [1–3, 6, 7]. Recent evidence, however, suggests that phenotypic state also correlates with the composition of the symbiotic bacterial community [8, 9].

Host-associated microbial communities are fundamental to the metabolism, immunity, and development of the host [10–12]. Symbiotic relationship between host and microbiota have deep evolutionary origins, often exhibit co-divergence over evolutionary time, and can be important for coping with environmental stressors [13, 14]. When faced with heterogeneous environments that are known to induce phenotypic plasticity, an animal may recruit, expel, and/or shuffle the membership and relative proportion of associated microbiota to assemble a community with specific functional properties (*e.g.*, genes and metabolic pathways) for the environmental conditions [12, 15–18]. When faced with prolonged diet restriction, for example, the composition and diversity of invertebrate- and vertebrate-associated bacterial communities shifts considerably to buffer against reduced exogenous nutrients [9, 19].

Planktotrophic (feeding) larvae of benthic marine inhabit coastal seas, where the abundance and distribution of exogenous nutrients are spatially and temporally heterogeneous [20]. Several groups of planktotrophic larvae, including sea urchins (phylum Echinodermata), respond to heterogeneous feeding environments by exhibiting morphological plasticity [21]. To increase individual fitness in the face of diet-restriction, larval sea urchins suppress development of the larval body and absorb stomach tissues to allocate energetic resources towards the feeding structures that, in turn, increases the capacity to collect particulates [22–27]. Moreover, morphological plasticity by echinoid larvae is coupled with physiological plasticity [28, 29], whereby diet-restriction correlates with reduced expression of genes associated with growth and metabolism and higher expression of genes involved with neurogenesis, environmental sensing, immunity, and longevity [29].

Larval sea urchins associate with bacterial communities that are diverse and dynamic yet specific to host species [9, 30], variable between populations [31], and distinct from the environmental microbiota [9, 30, 31]. Larval-associated bacterial communities shift over the course of embryonic development [30], with disease [32], and in response to food availability [9, 31, 33]. Specifically, the bacterial community associated with echinoid larvae exhibits bi-directional shifts that correlate with phenotype [9]. This correlation between morphological plasticity and the composition of the associated bacterial community, thus far, is unique to echinoid larvae, but recent data suggests similar responses in taxa from disparate clades [8]. What remains unclear is whether associating with a phenotype-specific bacterial community occurs before, during, or following the expression of morphological plasticity.

The expression of morphological plasticity by temperate Strongylocentrotids requires a multi-week stimulus [9, 22, 23, 31], which is restrictive for defining the temporal succession by the holobiont during morphological plasticity. Tropical and subtropical sea urchins, on the other hand, develop at a relatively rapid pace and may express plasticity in a few days [34–36]. This shortened developmental window permits a narrow time frame for changes in morphology and associations with bacteria that can be studied at a finer grain than species with prolonged larval durations. Using *Lytechinus variegatus*, a subtropical/tropical sea urchin known to exhibit morphological plasticity in four days [35–37], we define this temporal succession and test the hypothesis that shifts in the bacterial taxa precede a morphological response. To test this, we performed a differential feeding experiment on *L. variegatus* larvae and used amplicon sequencing to profile the larval-associated bacterial communities over the course of morphological plasticity.

## 2. Materials and methods

### (a) Specimen collection and larval rearing

Adult *L. variegatus* were collected from populations in Back Sound, North Carolina (NC), USA in July 2017, were transferred to the Duke University Marine Laboratory (Beaufort, NC) within one hour, and were maintained in flow-through aquaria.

Within two days of collections adult sea urchins were spawned by a one to two mL intracoelomic injection of 0.50 M potassium chloride. Gametes from five males and five females were pooled separately and the fertilization of eggs (diameter: 97.9 µm ± 1.07 µm) and larval rearing followed Strathmann [38], with the exception that embryos and larvae were reared using 5.0 μm filtered seawater (FSW) to include the environmental microbiota. Briefly, eggs were fertilized (verified using stereomicroscope) using dilute sperm in 100 mL of FSW at ambient temperature and salinity (Figure S1). After fertilization, embryos were transferred to jars with 3 L of FSW (n=4; Table S1) and diluted to a density of eight individuals•mL^−1^.

For larval feeding, monocultures of the cryptophyte *Rhodomonas lens* were grown in f/2 media at room temperature with a combination of ambient and artificial lighting for 24 hours per day [39].

### (b) Experimental feeding and tissue collection

At 24 hours post-fertilization, each jar of prism stage larvae was sub-divided into four replicates (n=16), diluted to two individuals•mL^−1^ in 3 L of FSW, and, at random, were provided growth medium-free *R. lens* at either 10,000, 1,000, 100, or 0 cells•mL^−1^ (n=4 per diet; Table S1). For the rest of the experiment, larval cultures had daily water changes of 90-95% volume and *R. lens* was replenished to the experimental density.

Larvae were differentially fed for five days and samples (n=100 larvae) were taken daily for each replicate of the four treatments. Immediately after sampling, larval samples were concentrated into a pellet using a microcentrifuge and the FSW was removed with a sterile glass pipette. Pelleted larvae were then preserved in RNAlater (Thermo Scientific, Massachusetts, USA) and stored at −20°C until extraction of nucleic acids.

### (c) Larval morphology

In addition to sampling the larval-associated bacterial community, twenty larvae (n=20) from a single replicate of each dietary treatment were sampled for morphometric analysis. Larvae were imaged using a compound microscope (Leitz Labovert; camera: Olympus DP71) and morphometrics (length of larval body, MBL; post-oral arms, POA; and stomach area; Figure 1) were performed using ImageJ (v. 1.9.2; [40]). Statistical differences in larval morphology and stomach volume were compared using a two-way analysis of variance (ANOVA, cut off p=0.05) in JMP Pro (v. 13) to test whether larval morphology differed across time and between diets. When statistical differences were detected, we then performed a Tukey’s post-hoc test for pairwise comparisons.

**Figure 1.**
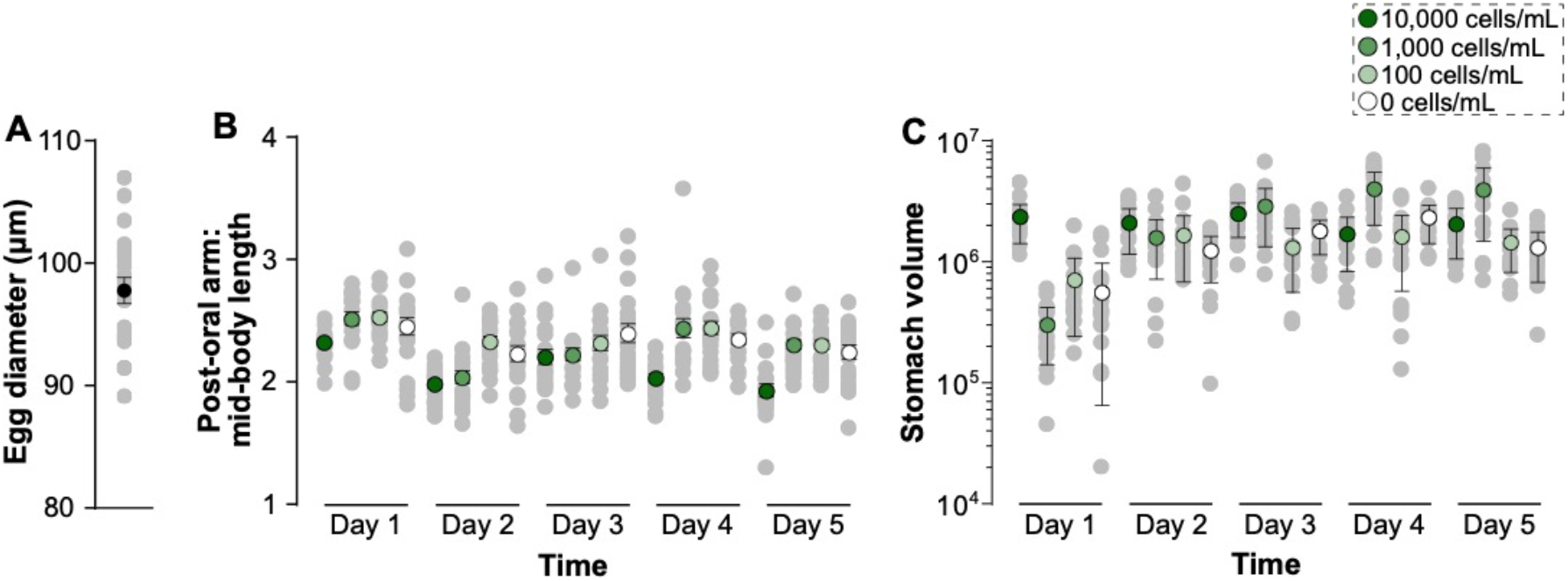
Egg size and morphometrics of *Lytechinus variegatus* larvae. (A) Mean egg diameter (± standard error) of unfertilized eggs. (B) Post-oral arm to mid body line ratio and (C) stomach volume (± standard error) of larvae having been fed either 10,000 (dark green), 1,000 (medium green), 100 (light green), and 0 cells•mL^−1^ (white) over the course of five days.

### (d) Assaying bacterial communities

Total DNA was extracted from larval samples using the GeneJet Genomic DNA Purification Kit (Thermo Scientific), quantified using the NanoDrop 2000 UV-Vis Spectrophotometer (Thermo Scientific), and diluted to 5 ng•μL^−1^ using RNase/DNase-free water.

Bacterial DNA was amplified using primers for the V3/V4 regions of the 16S rRNA gene (Table S2; [41]). Products were purified using the Axygen AxyPrep Mag PCR Clean-up Kit (Axygen Scientific), indexed via PCR using the Nextera XT Index Kit V2 (Illumina Inc.), and then purified again. At each of these three clean up states, fluorometric quantitation was performed using a Qubit (Life Technologies) and libraries were validated using a Bioanalyzer High Sensitivity DNA chip (Agilent Technologies). Illumina MiSeq sequencing (v3, 2×300 bp paired-end reads) was performed at the University of North Carolina at Charlotte. PCR recipe and thermal profiles are available in Table S2.

### (e) Computational analysis

Raw reads, along with quality information, were imported into QIIME 2 (v. 2019.1; [42]), where forward and reverse reads were paired using VSEARCH [43], filtered by quality score, and denoised using Deblur [44]. QIIME 2-generated ‘features’ were grouped into operational taxonomic units (OTUs) based on a minimum 99% similarity and were assigned taxonomy using SILVA (v. 132; [45]). Sequences matching to Archaea as well as samples with less than 1,000 reads were discarded and the filtered biom table was rarified to 1,287 sequences (*i.e.*, the read count for the sample with the least remaining reads; Figure S2).

To test whether community membership and composition shift over time, in response to food availability, and relative to morphology, we calculated unweighted and weighted UniFrac [46] values and compared them using principal coordinate analyses (PCoA). Results from these analyses were then recreated and stylized using QIIME 1 (v. 1.9.1; [47]) and Adobe Illustrator CC. To test for differences in membership and composition over the course of the differential feeding experiment, we used a two-way PERMANOVA and, subsequently, performed pairwise comparisons. To complement UniFrac values, we also calculated several measures of alpha diversity (*i.e.*, total OTUs, phylogenetic distance, McIntosh evenness, and McIntosh dominance) over time and across diets, and compared these values with a two-way analysis of variance (ANOVA) and, subsequently, performed by a Tukey’s post-hoc test for pairwise comparisons between diets and times. Lastly, we summarized the bacterial groups associated with *L. variegatus* larvae and determine which differ with diet and time using a two-way ANOVA.

The QIIME2 pipeline used to convert raw reads to OTUs for visualization of this data is presented in detail in Note S1.

## 3. Results

As a prefeeding *L. variegatus* larva, the associated bacterial community is comparably ‘simple’ and significantly different in membership and composition from all later larval stages (Figure 2; Tables S3-4). At the prefeeding stage, the associated bacterial community has less OTUs (~135 ± 5), is less phylogenetically diverse, and is less taxonomically dominant (*i.e.*, more even) than the bacterial community at each ensuing day (Figure 3A-B, Figure S3).

**Figure 2.**
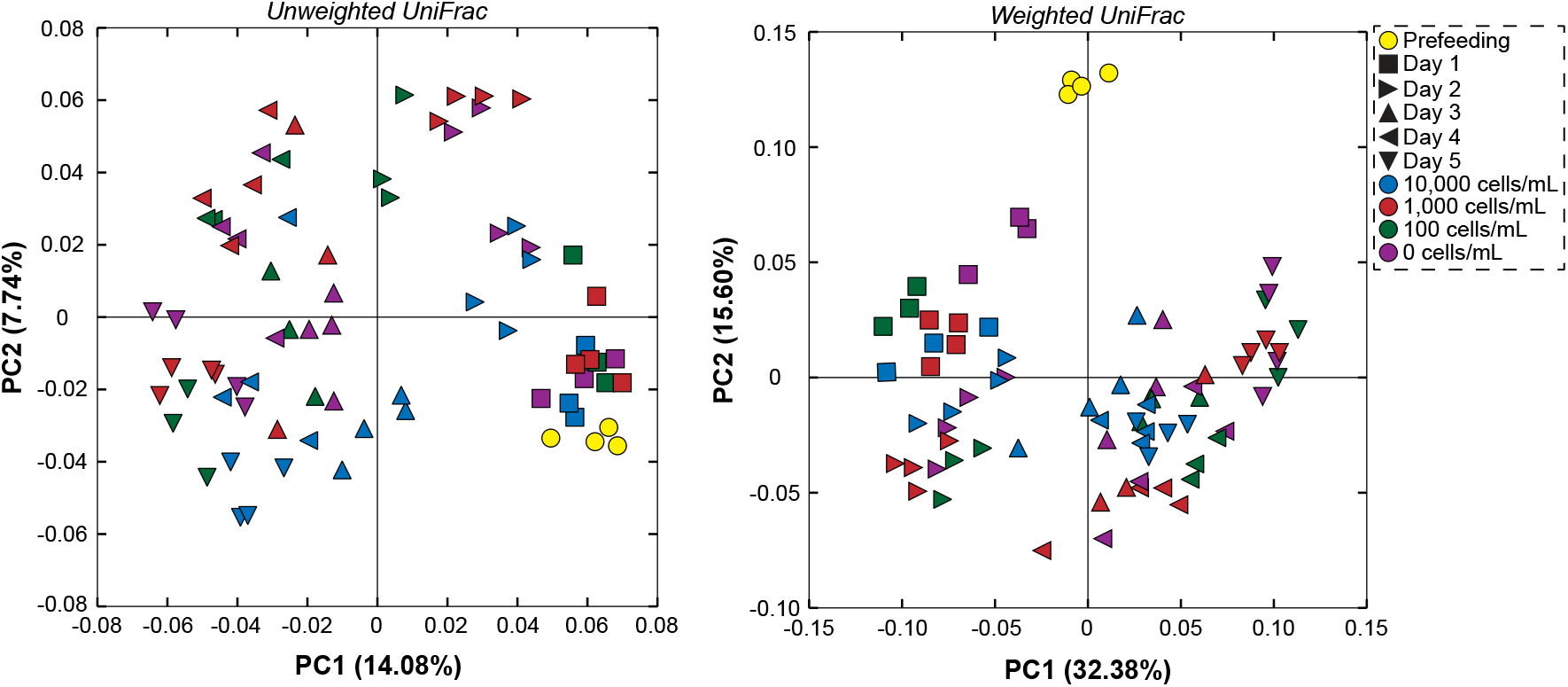
Dietary and temporal shifts in the bacterial community associated with *Lytechinus variegatus* larvae. Community similarity of *L. variegatus* larval-associated bacterial communities based on food availability (10,000, 1,000, 100, and 0 cells•mL^−1^ of a phytoplankton represented by blue, red, green and purple, respectively) over a multi-day exposure (prefeeding were yellow-circles while day 1, 2, 3, 4, and 5 were represented by a square, rightward triangle, upward triangle, leftward triangle, and downward triangle, respectively). Comparisons between food availability and over time are based on unweighted (left) and weighted (right) UniFrac values.

**Figure 3.**
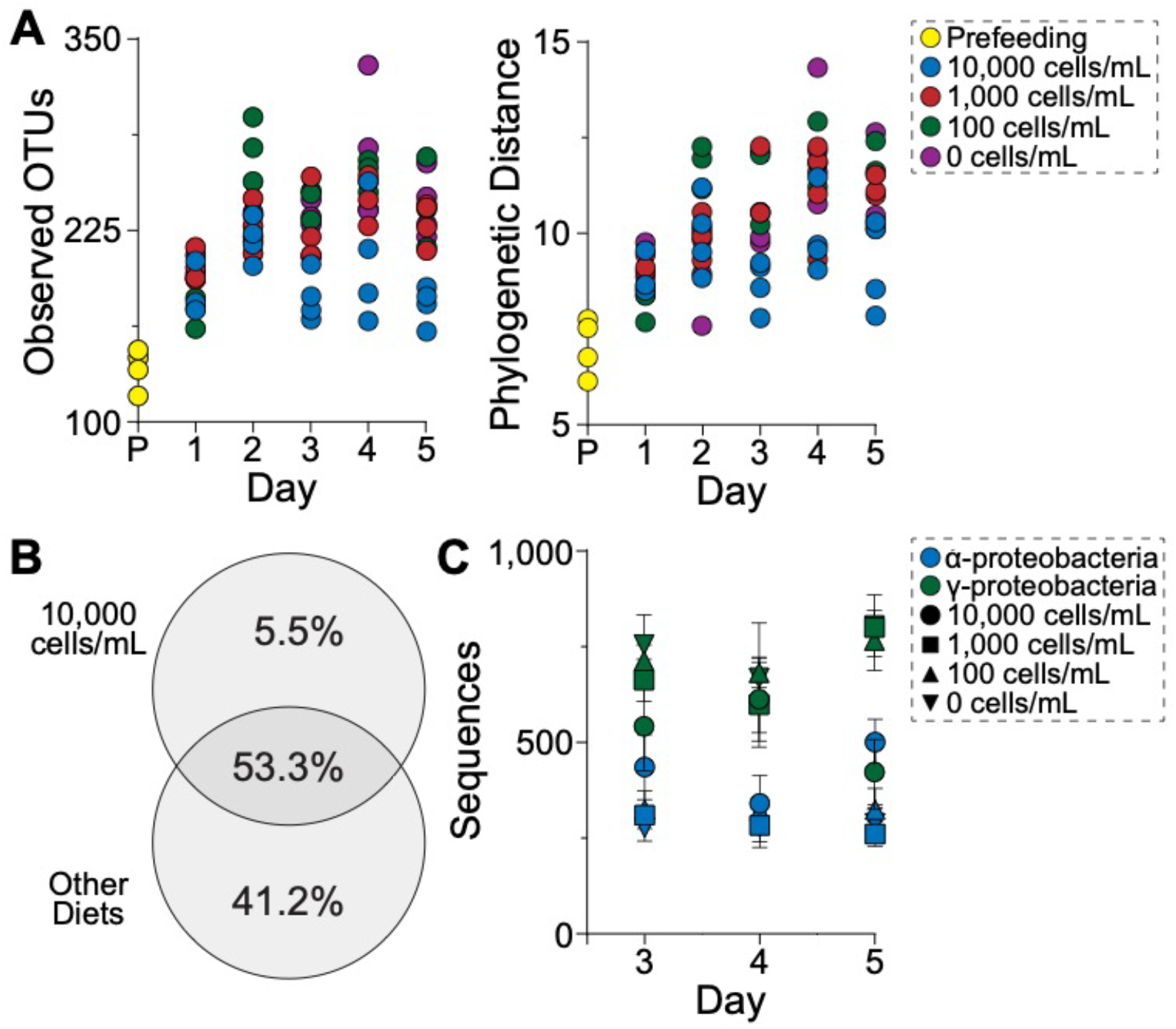
Structural changes to the bacterial community associated with *Lytechinus variegatus* larvae. (A) Enumeration of operational taxonomic units (OTUs) and phylogenetic distance of those OTUs for *L. variegatus* larvae from prefeeding through five days of feeding on either (10,000, 1,000, 100, and 0 cells•mL^−1^ of a phytoplankton represented by blue, red, green and purple, respectively). (B) Unweighted distribution of bacterial taxa for *L. variegatus* larvae from either of the larval phenotypes, and the (C) total sequences (from the rarefied table) of α- and γ-proteobacteria for these time points and diets.

Following one day of development and differential feeding, prefeeding larvae from each diet became two-arm larvae (Table S5), with the POA:MBL of larvae fed 10,000 and 0 cells•mL^−1^ was significantly lower than those fed 1,000 and 100 cells•mL^−1^ (Figure 1B; Tables S6). The membership, composition, phylogenetic diversity, and community evenness of the larval-associated bacterial community was not significantly different across feeding regimes while total OTUs and community dominance varied slightly between dietary treatments (Figures 2, 3, S3; Table S7).

A second day of differential feeding resulted in more rapid development for larvae in the highest food concentration. Larvae fed 10,000 cells•mL^−1^ were four-arm larvae while those fed 1,000, 100, and 0 cells•mL^−1^ were maintained as two-arm larvae (Tables S5). Despite this difference in developmental stage, larvae fed 10,000 and 1,000 cells•mL^−1^ had a similar POA:MBL, which was lower than those fed 100 and 0 cells•mL^−1^ (Figures 1; Tables S6). Dietary treatments also resulted in differences in bacterial communities, where algal concentrations induced diet-specific bacterial communities in both membership and composition (Figure 2; Tables S3-4). This led to larval-associated communities with more OTUs and a higher phylogenetic diversity but similar community evenness and dominance (Figure 3A-B, S3; Table S7), with larvae fed 100 cells•mL^−1^ being the most diverse and those fed 10,000 and 1,000 cells•mL^−1^ being the least diverse (Figure 3A-B, S3; Table S7).

Independent of diet quantity, *L. variegatus* larvae fed at each algal concentration for three days were all four-armed with statistically similar POA:MBL (Figure 1; Table S5, S6). The membership and composition of the bacterial communities associated with larvae fed 1,000, 100, and 0 cells•mL^−1^ were similar while those fed 10,000 cells•mL^−1^ differed significantly (Figure 2; Tables S3-4). A reduction in total OTUs and phylogenetic distance as well as a more even and less taxonomically dominant community resulted in clear differential structuring in the bacterial community for larvae fed 10,000 cells•mL^−1^ (Figures 3, S3; Table S7). This response was not observed for larvae fed 1,000, 100, and 0 cells•mL^−1^, as these treatments maintained their community structure from the day prior (Figures 3, S3; Table S8).

As expected based on previous literature for *L. variegatus* [35, 36], morphological plasticity was observed on the fourth day of differential feeding (Figure 1; Tables S5, S6), where larvae fed 1,000, 100, and 0 cells•mL^−1^ had a higher POA:MBL than those fed 10,000 cells•mL^−1^. Like the day prior, larvae fed 1,000, 100, and 0 cells•mL^−1^ associated with bacterial communities that were similar in membership and composition while larvae fed 10,000 cells•mL^−1^ differed significantly (Figure 2; Tables S3-4). As morphological plasticity was expressed, the number of OTUs for larvae in these three lower food concentrations had similar phylogenetic distance (Figure 3; Table S7); however, larvae fed 10,000 cells•mL^−1^ continued to associate with a comparatively more even and less dominant community.

Consistent with the previous day, *L. variegatus* maintained expression of morphological plasticity (Figure 1; Table S6), a result that was not confounded by a continuation in larval developmental stage (Table S5). On day five of differential feeding, patterns of community membership and composition (Figure 2; Tables S3-4) as well as total OTUs, phylogenetic distance, evenness, and dominance all reflected the differences observed in larval morphology (Figure 3; Table S7). Specifically, larvae fed 10,000 cells•mL^−1^ associated with less OTUs that span less phylogenetic distance while maintaining a more even and less dominant community than larvae fed 1,000, 100, and 0 cells•mL^−1^ (Figure 3; Table S7).

Spanning the induction and expression of morphological plasticity, prefeeding and feeding larval stages of *L. variegatus* consisted primarily of three Bacteroidetes (23.5%) families and eleven Proteobacteria (64.0%) families (Figure S4). Over the three days where morphological plasticity was expressed, we observed the abundance of all 14 bacterial families from Bacteroidetes and Proteobacteria differ significantly with time while 11 (of 14) of these bacterial families differed across diets (Table S9). Moreover, we observed that the abundance of α- and γ-proteobacteria were inversely proportional (Figure 3C, S4; Table S10), such that α-proteobacteria increased following plasticity while γ-proteobacteria decreased.

Across the same three days where morphological plasticity was observed, larvae fed 10,000 cells•mL^−1^ stably associated with 114 OTUs (43.7%) while 16-40 OTUs (6.1-17.6%) were specific to a single day and 19-34 OTUs (7.3-13.0%) were shared between successive days (Figure 3). Larvae fed 1,000, 100, and 0 cells•mL^−1^, on the other hand, stably associated with an average of 190 OTUs (56.6%) while 15-19 OTUs (4.5-5.7%) were specific to a single day and 36-39 OTUs (10.7-11.6%) were shared between successive days (Figure 3). Despite the shifts in community membership, the larval stages of *L. variegatus* maintained a ‘core’ of 31 OTUs from the Bacteroidetes (12.9%), Epsilonbacteraeota (3.2%), and Proteobacteria (83.9%).

## 4. Discussion

Morphological plasticity is an adaptive response to heterogenous environments when a fitness advantage results relative to an individual with no response [1–3]. Larval sea urchins, in particular, increase individual fitness in dilute feeding environments by elongating their feeding structure that, in turn, allows for a greater capacity to filter particulates and reduce development time [21–23, 26, 34, 36]. Morphological plasticity for larval sea urchins is also coupled with significant shifts in the associated bacterial community [9, 31]. The timing of morphological plasticity and associating with a phenotype-specific bacterial community, however, remains unclear.

Daily profiling of the bacterial communities associated with *L. variegatus* larvae over the course of early development and through morphological plasticity supports three primary findings. First, a four-stage successional pattern is followed as larvae transition towards phenotype-specific bacterial communities. Second, the relatedness of the larval-associated bacterial community directly reflects morphological plasticity and does so prior to the expression of the phenotype. Third, relatedness of the bacterial communities prior to and following morphological plasticity implies that the long-arm is the default phenotype.

From the initiation of differential feeding through the expression of morphological plasticity and associating with phenotype-specific bacterial communities, we observed four specific stages for *L. variegatus* larvae. First, ‘Stage 1’ followed one-day of differential feeding where the bacterial communities across diets were similar in membership and composition (Figure 4). Second, ‘Stage 2’ followed two-days of differential feeding where the membership and composition of the larval-associated bacterial communities were diet-specific (Figure 4). Third, ‘Stage 3’ was observed after three-days of differential feeding where the membership and composition of the larval-associated bacterial communities reflected the larval phenotypes while the host had yet to (Figure 4). Lastly, ‘Stage 4’ was observed following four- and five-days of differential feeding and was where the relatedness of the bacterial communities correlated with morphological plasticity (Figure 4).

**Figure 4.**
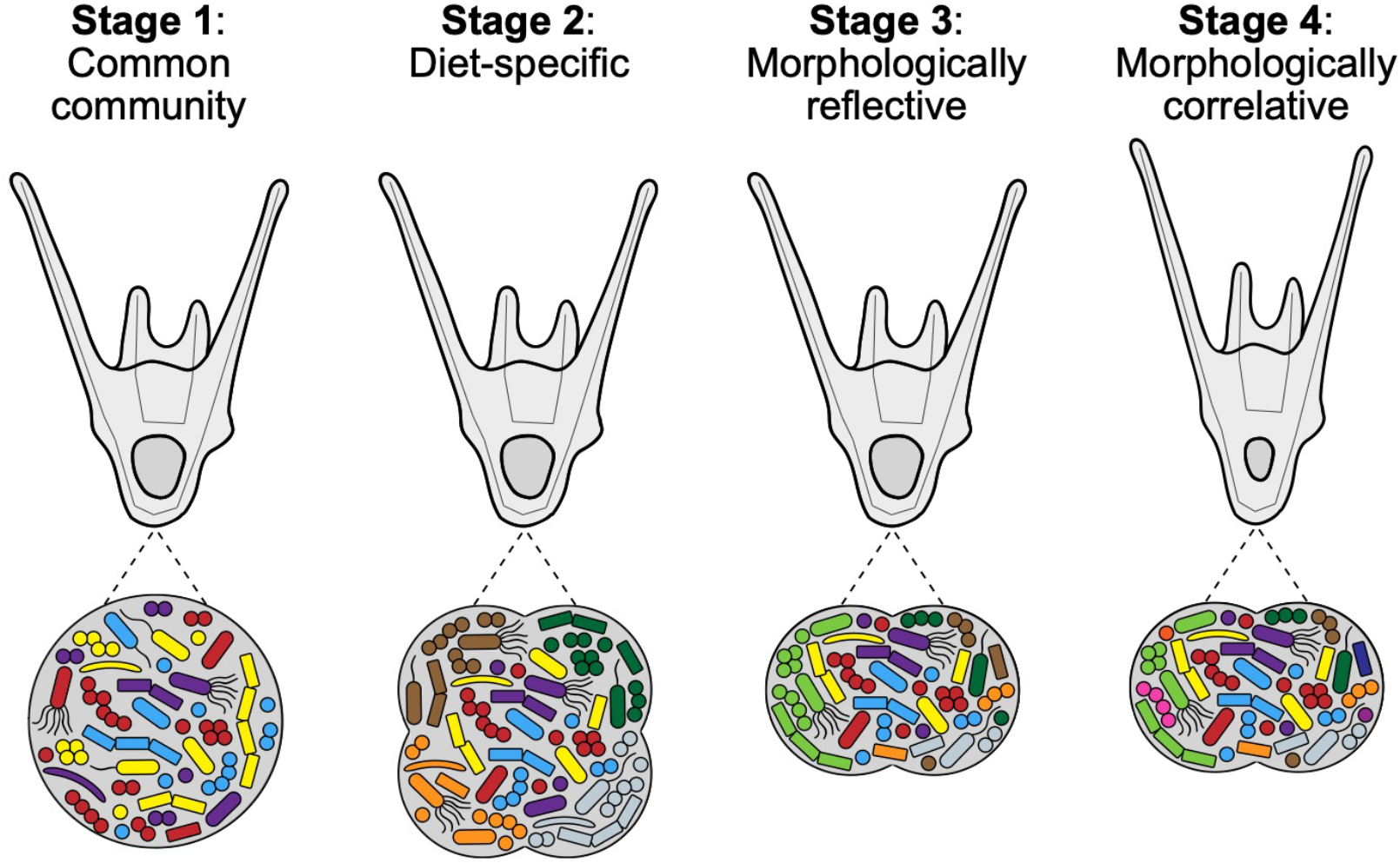
Dynamics of the bacterial community during morphological plasticity. Visual model representation of morphological plasticity for echinoid larvae and the four successive stages towards associations with a phenotype-specific bacterial community.

The expression of morphological plasticity by temperate Strongylocentrotids [9] show similar ‘Stages’ to the patterns observed with subtropical/tropical *L. variegatus* larvae. Specifically, over four weeks of differential feeding for Strongylocentrotids, it was observed that these larvae exhibit ‘Stages 1, 2, and 4’ [9]. The temporal pattern of these ‘Stages’ was, however, unclear. Timing by *L. variegatus* larvae provides a fine grain temporal organization to compare with Strongylocentrotid larvae as well as implies another stage that was potentially missed. Similarly, differential feeding of a larval sea star (*Acanthaster* sp.) showed both ‘Stages 1 and 2’ [33], which suggests that these stages may be broadly conserved in these two distinct classes (Echinoidea and Asteroidea). Consistent observation of these stages in the literature may imply that these four stages (Figure 4) are common for echinoderm larvae. Whether these ‘Stages’ are observed for other taxonomic Classes known to also exhibit morphological plasticity (*e.g.*, Ophiuroidea [48] and Holothuroidea [49]) is unknown but merits testing.

Time lags between experiencing and responding to an environmental stimulus are common, advantageous, and hypothesized to be under strong selection for species expressing morphological plasticity [4, 5]. For plasticity to be advantageous, time lags must be short relative to the time scale of the environmental variant but long enough to minimize false-positive expression [4, 5, 50]. When faced with fine grain variability in food abundance, *L. variegatus* larvae are unable to match phenotype with feeding environment, as the lag for morphological responses exceeds two days [51]. Despite a multi-day time lag, *L. variegatus* larvae are capable of associating with bacterial communities reflective of morphological plasticity prior to expressing the morphological trait. This, in principle, implies that instances where a time lag exceeds the morphological response (*e.g.*, [51]), a holobiont may exhibit shifts in the associated bacterial community corresponding with the environmental variation in absence or anticipation of morphological plasticity. The expression of morphological plasticity also comes with the inherent energetic cost of producing and maintaining an alternate phenotype [7]; thus, shifts in the bacterial community may be energetically favored over morphological plasticity in heterogenous environments.

Provided the substantial energetic requirement for planktotrophic echinoids to undergo larval development [52], laboratory culturing has traditionally simulated conditions equivalent to phytoplankton blooms. This, in turn, has instilled the general idea that morphological plasticity requires food deprivation and plasticity is the elongation of the feeding apparatus [21, 23, 35]. Field culturing, however, suggests the opposite, that echinoid larvae are naturally food-limited [20, 53, 54] and ‘plasticity’ is shortening the larval arm and expanding stomach volume in response to uncommon food-rich environments [23]. The molecular mechanisms underlying morphological plasticity have suggested that algal chemosensations inhibit growth of the larval arms [28] and, thus, the long-arm phenotype would likely be the default under natural conditions.

Following the expression of morphological plasticity, the membership, composition, and structure of the bacterial communities associated with long-arm *L. variegatus* larvae mirrored larval siblings prior to plasticity. Simultaneously, the membership and composition of the bacterial communities associated with short-arm *L. variegatus* larvae differed significantly from both pre-plasticity and long-arm siblings. This community shift followed a reduction in total and short-arm-specific OTUs and phylogenetic diversity of a more even and less dominant community. This implies that well-fed (or short-arm) larvae deviate from the microbial trajectory of larvae having yet to express plasticity and that the associated bacterial community may play a role in regulating the short-arm phenotype.

Taken together, the data presented here suggests: (*i*) that phenotype-specific bacterial communities for larval *L. variegatus* follow a four-stage progression (Figure 4); (*ii*) that shift in bacterial taxa precedes morphological plasticity and occur during the time lag; and (*iii*) that the long-arm phenotype is most similar to pre-plasticity larval siblings and the short-arm phenotype correlates with a restructuring of the bacterial community. Determining if and how this community contributes to larval fitness during before, during, and following the expression of morphological plasticity merits future investigation and would require multi-omic comparisons [55] between axenic and germ-rich siblings (*e.g.*, [56–58]) as well as add-back experiments of individual bacterial taxa (*e.g.*, [59–61]).

## Acknowledgements

We thank Daniel Rittschof (Duke Univ.) for providing laboratory space; Beatriz Orihuela (Duke Univ.) for endless logistical assistance; Josh Osterberg (Duke Univ.) for collecting adult urchins; Karen Lopez (UNC Charlotte) for technical assistance with sequencing; Daniel Janies (UNC Charlotte) for sequencing resources; and Justin McAlister (Holy Cross Univ.) and Jason Hodin (Univ. Washington) for discussions on morphological plasticity in echinoid larvae.

## Data accessibility

The raw sequence reads as part of this dataset are available on the Dryad Digital Repository.

---

**Figure S1.**
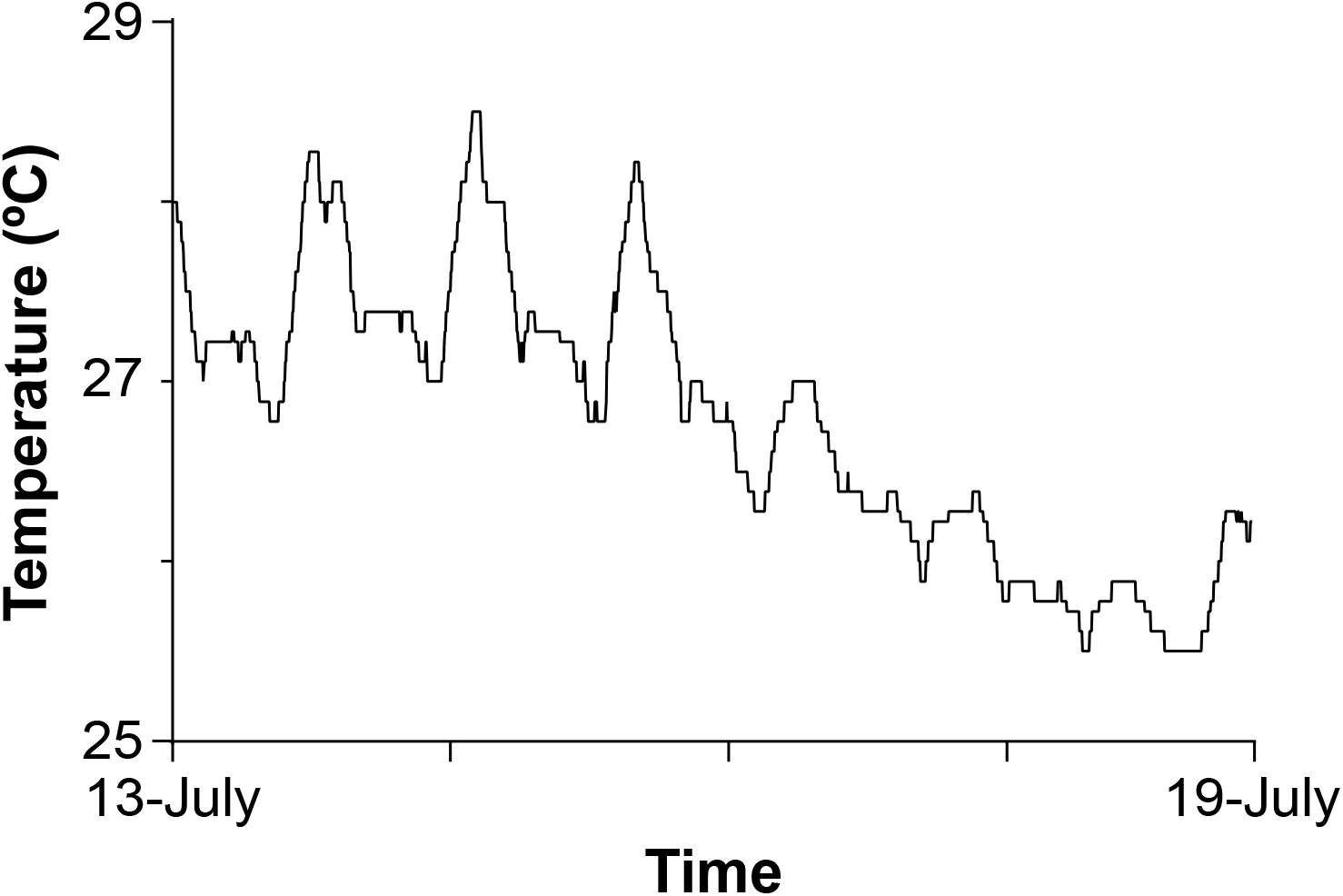
*in situ* measurements of sea surface temperature at the Duke University Marine Laboratory. Sea surface temperature at Beaufort, NC (NOAA station, BFTN7) for the entirety of larval experimentation (13-19 July 2017).

**Figure S2.**
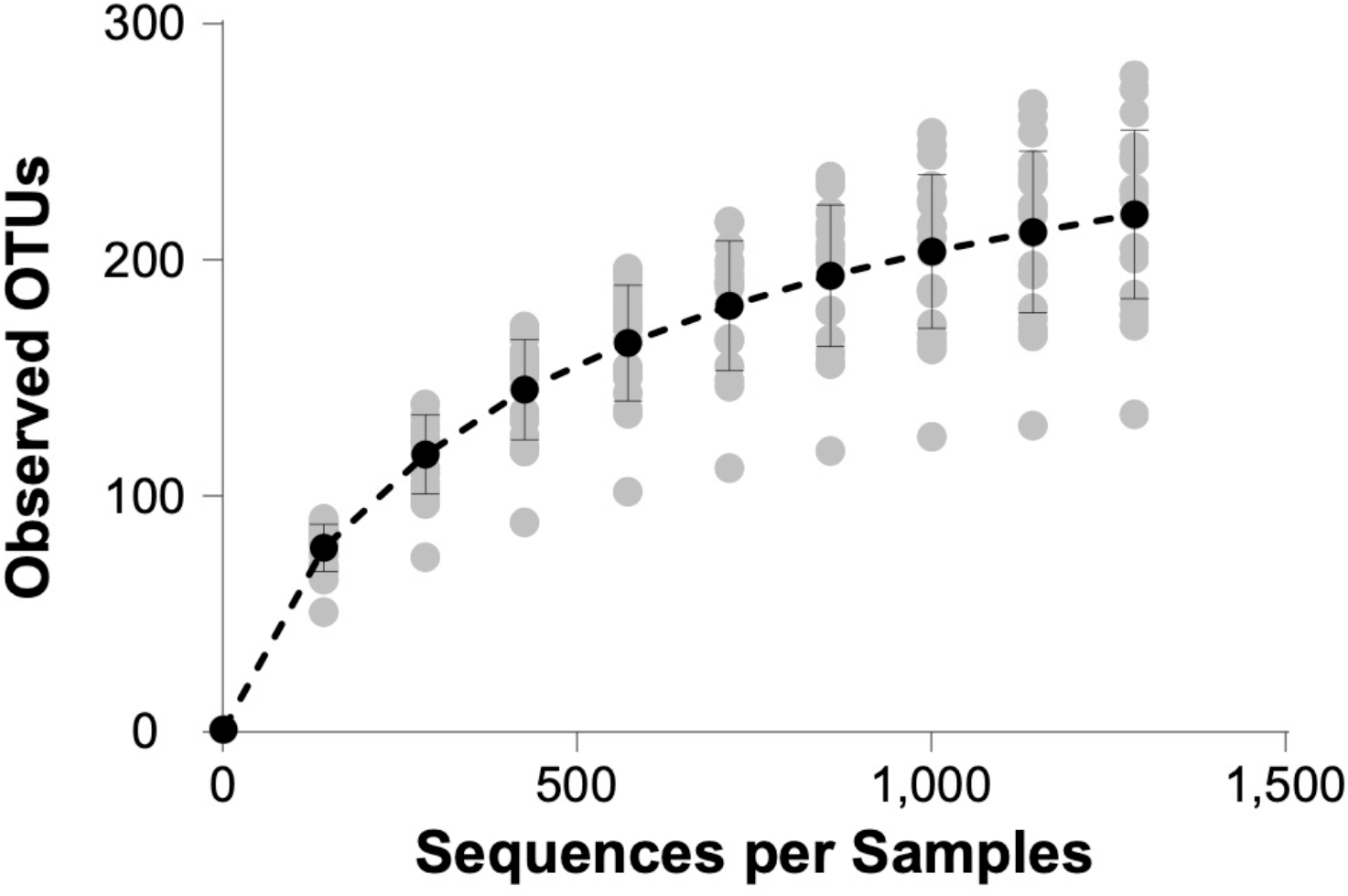
Alpha rarefaction curve for *Lytechinus variegatus* larvae. Alpha rarefaction curve for the associated bacteria with for *L. variegatus* from prefeeding through five days of feeding on either 10,000, 1,000, 100, and 0 cells•mL^−1^ of a phytoplankton based on the rarefaction depth (1,287 sequences) used for all analyses.

**Figure S3.**
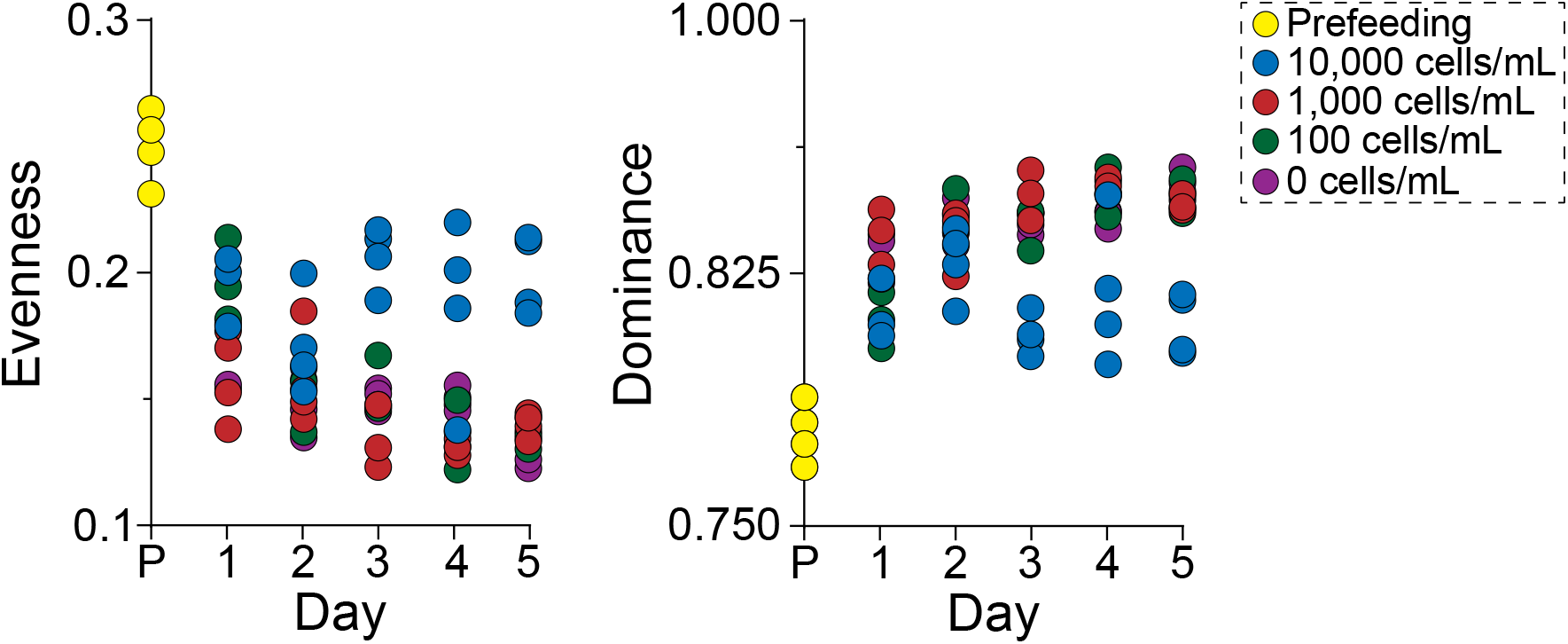
Changes in alpha diversity metrics to the bacterial community associated with *Lytechinus variegatus* larvae. Evenness and dominance of bacterial taxa for *L. variegatus* larvae from prefeeding through five days of feeding on either 10,000, 1,000, 100, and 0 cells•mL^−1^ of a phytoplankton (represented by blue, red, green and purple, respectively).

**Figure S4.**
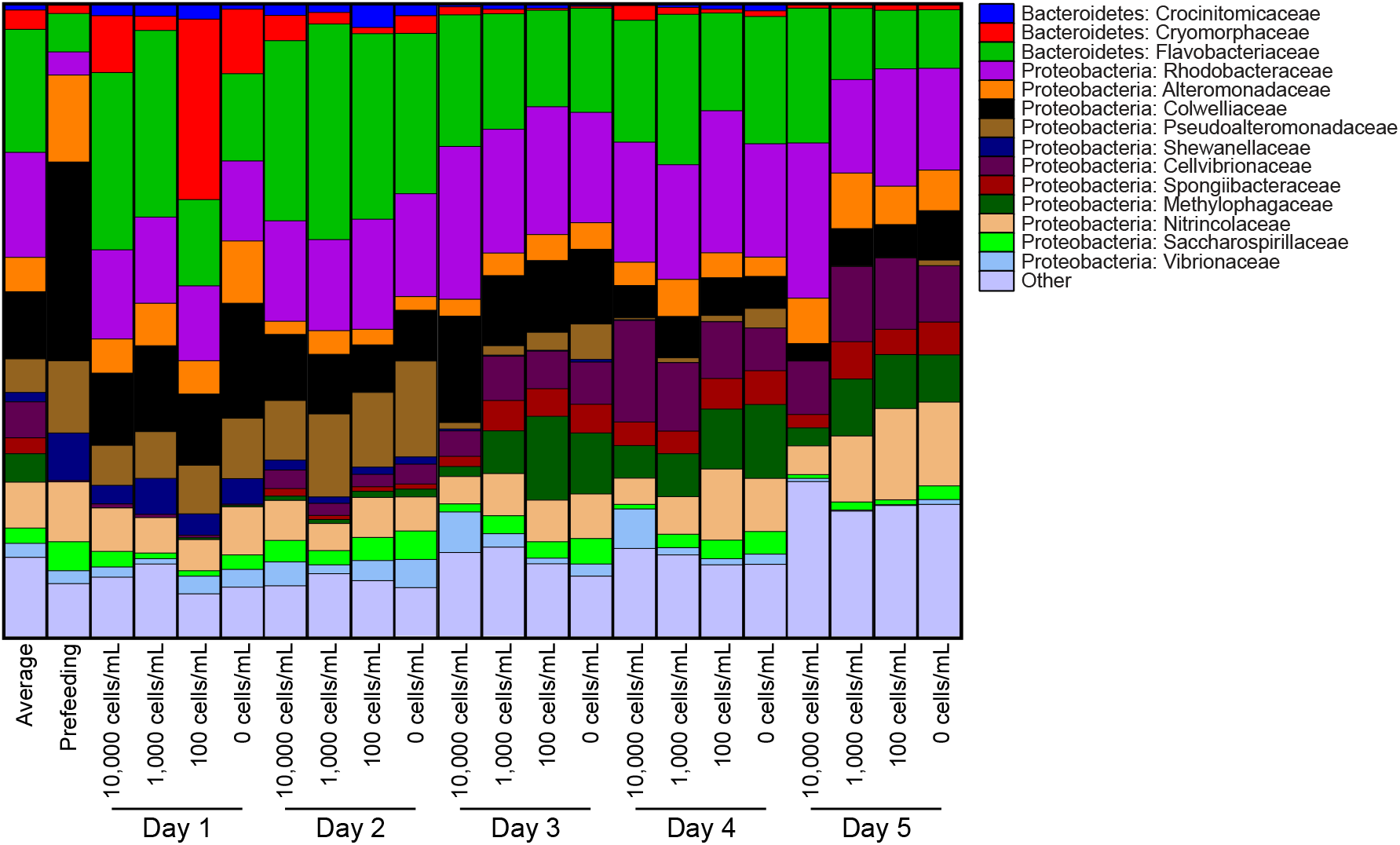
Bacterial taxa associated with differentially fed *Lytechinus variegatus* larvae. Bacterial family associated with *L. variegatus* larvae fed either fed either 10,000, 1,000, 100, or 0 cells mL^−1^ over the course of five days of differential feeding.

